# Molecular consequences of *PQBP1* deficiency, involved in the X-linked Renpenning syndrome

**DOI:** 10.1101/2022.05.29.493091

**Authors:** Jérémie Courraud, Camille Engel, Angélique Quartier, Nathalie Drouot, Ursula Houessou, Damien Plassard, Arthur Sorlin, Elise Brischoux-Boucher, Lionel Van Maldergem, Evan Gouy, Massimiliano Rossi, Patrick Edery, Audrey Putoux, Brigitte Gilbert-Dussardier, Vera Kalscheuer, Jean-Louis Mandel, Amélie Piton

## Abstract

Mutations in the *PQBP1* gene (polyglutamine-binding protein 1) are responsible for a syndromic X-linked form of intellectual disability (XLID), the Renpenning syndrome. *PQBP1* encodes a protein that plays a role in the regulation of gene expression, splicing and mRNA translation. To investigate the consequences of variants in *PQBP1*, we performed transcriptomic studies in 1) patients’ lymphoblastoid cell lines (LCL) carrying pathogenic variants in *PQBP1* and 2) in human neural stem cells (hNSC) knocked-down (KD) for *PQBP1*. This led to the identification of a hundred dysregulated genes. In particular, we identified an increase in the expression of a non-canonical isoform of another XLID gene, *UPF3B. UPF3B* plays a crucial role during neurodevelopment by coding for an important actor of the nonsense mRNA mediated decay (NMD) system involved in regulation of protein translation, however, the exact function of the non-canonical isoform,*UPF3B_S*, is currently unknown. In order to investigate the role of *UPF3B_S* isoform, we compared the protein interactome of UPF3B_S to the canonical isoform (UPF3B_L). We confirmed that, on the contrary to UPF3B_L, UPF3B_S does not interact with the UPF2/UPF1 complex while it still interacts with exon junction complexes (EJC). However, no notable decrease of NMD pathways was observed in patient’s LCL or in hNSC KD for *PQBP1*. We identified several additional protein interactors specific to UPF3B_S. Moreover, we used the increase of *UPF3B_S* mRNA as a molecular marker to test the pathogenicity of variants of unknown clinical significance identified in individuals with ID in *PQPB1*. We analyzed patients’ LCL mRNA as well as blood mRNA samples and performed complementation studies in HeLa cells by overexpressing Wild-type and mutant *PQBP1* cDNA. We showed that all these three approaches were efficient to test the effect of variants, at least for variants affecting the CTD domain of the protein. In conclusion, our study provides information on how *PQBP1* deficiency may affect the expression of genes and isoforms, such as *UPF3B*. This informs about the pathological mechanisms involved in Renpenning syndrome but also allows to propose a functional test for variants of unknown significance identified in *PQBP1*.

## INTRODUCTION

More than a dozen of pathogenic variants, mostly truncating, have been described in *PQBP1* (polyglutamine-binding protein 1), responsible for syndromic X-linked intellectual disabilities (XLID) (Renpenning, Sutherland□Haan, Hamel, Porteous, and Golabi□Ito□Hall) (1–4) regrouped now under the unique term of Renpenning syndrome (5). Clinical manifestations associated with these syndromes include microcephaly, growth retardation, lean profile and specific facial features (6).

*PQBP1* encodes a protein involved in different cellular processes such as regulation of transcription, splicing, translation or even response to retroviral infection. PQBP1 was initially described as a protein interacting with polyglutamine tracts of huntingtin or ataxin1, through its polar amino-acid-rich domain (PRD)(7). It also contains on its N-terminal side a WW domain which has a transcriptional activity and interacts with the splicing factor SIPP1/WP11. The C-terminal part of PQBP1 includes a domain (CTD) interacting with other splicing factors such as U5-15kDa/TXNL4A (8). Moreover, a nuclear localization signal (NLS) allows its addressing to the nucleus through the Kapβ2 receptor (9,10). In the nucleus, PQBP1 is involved in transcription regulation through its interaction with activated RNA polymerase II (7) and various transcription factors such as POU3F2/Brn2 (11). PQBP1 has been found to be located in nuclear speckles, suggesting a role in splicing regulation, consistent with its interactions with the different splicing factors cited above and with the general splicing alterations observed after *PQBP1* knock-down in a model of murine primary neurons (12). PQBP1 is a nuclear-cytoplasmic shuttling protein playing also various roles in the cytoplasm. Indeed, it can be localized in stress granules (13) and was shown to play a role in translation of messenger RNAs (14). PQBP1 binds to the elongation factor eEF2 via its WW domain and suppresses its phosphorylation-mediated inactivation. Loss of *PQBP1* leads to an increase of the phosphorylation of eEF2 stopping translational elongation and resulting in a global decrease of protein synthesis, which affect protein synthesis-dependent synaptic plasticity in the hippocampus (14). PQBP1 is involved in the regulation of neuronal ciliogenesis via its interaction with Dynamin2 (15). Finally PQBP1 can also play a role in response to retroviral infection, interacting with reverse-transcribed HIV-1 DNA, probably through its CTD domain, and to the viral DNA sensor cGAS probably through its WW domain, and therefore contributing to the innate immune response (16).

Constitutive knock-out of *Pqbp1* in mice appears to be lethal (www.mousephenotype.org/), mice models where *Pqbp1* was knocked-down (~50%) display abnormal anxiety-related behavior, and a decrease in anxiety-related cognition (17). Nestin-Cre conditional knock-out (*Pqbp1* cKO) display abnormal anxiety-related behavior, as well as abnormal fear conditioning and motor dysfunction at the rotarod test (18). Recently, the same group demonstrated that cKO mice showed a short stature and a reduction in bone mass, and display impairment in bone formation and chondrocyte deficiency with reduced osteoblast and chondrocyte-related gene expression (19). The *Pqbp1* cKO showed microcephaly, probably resulting from an elongated cell cycle of neural progenitors (18).

In humans, most of the variants described are small indels, located in the AG hexamer of exon 4 or downstream, resulting in truncated proteins lacking their C-terminal domain, but few additional truncating variants were also reported (6,20–22). Only few missense variants have been reported in *PQBP1* and for years, only a unique missense variant (p.Tyr65Cys) was considered as pathogenic (4). This missense variant affects a very conserved amino acid position located in the WW domain and disrupts the interaction between PQBP1 and the splicing factor WBP11 leading to a decreased pre-mRNA splicing efficiency (23). Two other missense variations (p.Arg243Trp and p.Pro244Leu), located in the CTD domain were identified more recently in patients with ID by our group and others (24,25). The Pro244Leu change was shown to disrupt PQBP1 binding to the splicing factor U5-15kDa/TXNL4A (10).

We describe in this study the changes in gene expression induced by a loss of function of *PQBP1* in human cells, using both lymphoblastoid cell lines (LCL) of patients carrying pathogenic variants in *PQBP1* and human neural stem cells (hNSC) where *PQBP1* was knocked-down (KD) by siRNA. We report in particular the increase of expression of a non-canonical isoform of the *UPF3B* gene, another gene involved in XLID playing a role in nonsense-mediated mRNA decay (NMD) and regulation of translation. Finally, we showed that we can use this increase as a biomarker for Renpenning syndrome to test the pathogenicity of variants of unknown significance in *PQBP1* located in the C terminal part of the protein.

## Material and methods

### Subjects

Clinical information was retrieved from clinicians for individuals carrying pathogenic variants identified by panel or exome sequencing. All *PQBP1* variants were named according to the isoform NM_005710.2 which encodes a protein of 265 amino acids. The potential effect of missense variants was predicted using PolyPhen2 (http://genetics.bwh.harvard.edu/pph2/) and CADD (https://cadd.gs.washington.edu/). LCL were available for individuals with pathogenic variants in *PQBP1* (n=7), with variants of uncertain significance (n=3), control individuals (n=6) and individuals with pathogenic variant in *FMRI* (Fragile X syndrome, FXS) (n=4) (**Figure S1, Table S1**). Blood RNA samples (Paxgene) were available for individuals with pathogenic variants (n=3) in *PQBP1*, VUS in *PQBP1* (n=1) pathogenic variants in *FMR1* (n=2) and individuals with other genetic forms of ID (n=5) (**Table S1**).

### Plasmids, siRNA and antibodies

A pool of 4 different siRNA directed against PQBP1 mRNA was used for *PQBP1* KD (1: UGACAGGGAUCGAGAGCGU; 2: AGCCAUGACAAGUCGGACA; 3: ACGAUGAUCCUGUGGACUA; 4: AGAUCAUUGCCGAGGACUA). For complementation test, PQBP1 inactivation was performed using siRNA number 4. Human *PQBP1* cDNA NM_005710.2 sequence (including UTR) was subcloned into PCS2+ plasmid under CMV promoter and optimized to escape to degradation by siRNA against *PQBP1* by introducing 3 silent point mutations. Benign, pathogenic and VUS variants were introduced by site-directed mutagenesis (**Table S1)**. Human *UPF3B_L* cDNA (ENST00000276201 or NM_080632.2) and *UPF3B_S* (ENST00000636792) were subcloned into pcDNA3 plasmid under CMV promoter and tagged with HA motif (YPYDVPDYA) at the C-terminal side of the protein. For western blots and immunostaining, specific primary antibodies were used: anti-UPF3B (NSJ BIOREAGENTS), anti-PQBP1 (NOVUSBIO), anti-APP (Invitrogen Invitrogen 51-2700), anti-GAPDH (G9545 Sigma Aldrich). The secondary antibodies used were HRP-labelled goat anti-mouse or anti-rabbit IgG and HRP (Jackson Immunoresearch, Baltimore, PA, USA).

### Cell culture, transfection and proliferation assay

Lymphoblastoid cell lines (LCL) were previously (1) or newly established by infection of blood lymphocytes using Epstein-Barr virus and were maintained in RPMI without HEPES and 10% fetal calf serum. Two lines of human neuronal stem cells (hNSC) were used: hNSC-1 (SA001) and hNSC-2 (GM01869), previously described (26,27). They were seeded on poly-ornithine and laminin-coated (Sigma-Aldrich) dishes and maintained in DMEM/F12 and Neurobasal medium (1:1) supplemented with N2, B27, 2-mercaptoethanol (Invitrogen, Carlsbad, CA, USA), BDNF (20ng/mL), FGF-2 (10ng/mL) (PeproTech, Rocky Hill, NJ, USA) and EGF (R&D Systems; 10ng/mL) as described (27). HEK293 and HeLa cells were maintained respectively in DMEM (1g glucose / liter), 10% fetal calf serum and gentamicin (40μg/mL) in CellBind^®^ flasks (Corning, NY, USA) for HEK293T, DMEM (1g glucose / liter), 5% fetal calf serum and gentamicin (40μg/mL) in CellBind^®^ flasks (Corning, NY, USA) for HeLa. Cells were transfected using INTERFERin (Polyplus) or Lipofectamine 2000 transfection protocol (Thermo Fisher Scientific) for siRNA and plasmids respectively. Cells were stopped at 24/48 and 96 hours after transfection for protein and RNA extractions. For proliferation assay, reverse transfection of SA001 and GM01869 hNSCs (6,400 cells/cm2) was performed in 96-well plates using INTERFERin reagent and 120nM siRNA according to manufacturer’s recommendation. Each day a plate was fixed and stained with DAPI and number of nuclei in each condition counted using CellInsight automated microscope and HCS Studio software (Thermo Fisher Scientific).

### mRNASeq

For the LCL cells, RNA-seq libraries of template molecules were prepared as previously described (27). For the hNSC, RNA-seq libraries of template molecules suitable for high throughput sequencing were established from 300ng of total RNA using the KAPA RNA HyperPrep Kit after a first step of purification using poly-T oligo-attached magnetic beads (Roche). Libraries were sequenced on the Illumina Hiseq 4000 sequencer as paired-end 100 base reads. Reads were mapped onto the hg38 assembly of the human genome using Tophat 2.0.10 or 2.0.14 (28) and the bowtie version 2-2.1.0 aligner (29) Gene expression was quantified using HTSeq-0.6.1 (30)), using intersection-nonempty mode and gene annotations from Ensembl release 85. Only non-ambiguously assigned reads were kept. Comparisons of interest were performed using R 3.2.5 and DESeq2 1.10.1 (31). More precisely, read counts were normalized across libraries with the median-of-ratios method proposed by Anders and Huber (30) and a Wald test was used to estimate the p-values. Adjustment for multiple testing was performed with the Benjamini and Hochberg method (31). Alternative splicing and isoform-specific differential expression were identified using DEXSeq 1.14.2 (32).

### RT-qPCR

Total RNAs were extracted from cells using the RNeasy extraction kit and treated with RNase free DNAse set during 20 min (Qiagen, Valencia, CA, USA) or from blood using PaxGene blood RNA kit extraction (Preanalytix, Hombrechtikon, Switzerland). RNA levels and quality were quantified using a Nanodrop spectrophotometer and then with a 2100 Bioanalyzer (Agilent, Santa Clara, CA, USA) for the RNA used for RNA-sequencing. For reverse transcription-PCR (RT-PCR), 500ng to 1μg of total RNA was reverse transcribed into cDNA using random hexamers and SuperScript IV reverse transcriptase according to manufacturer’s recommendation. Real-time PCR quantification (qPCR) was performed on LightCycler 480 II (Roche, Basel, Switzerland) using the QuantiTect SYBR Green PCR Master Mix (Qiagen). All qPCR reactions were performed in triplicate. Primer sets are listed in **Table S2**. Reaction specificity was controlled by post-amplification melting curve analysis. The relative expression of gene-of-interest *vs GAPDH* and *YWHAZ* was calculated using the 2-(ΔΔCt) method and a parametric Student’s t-test was performed in order to compare control vs patient cells or untreated vs siRNA treated cells. Error bars represent standard error of the mean (SEM).

### Western Blot

Cells were lysed in RIPA buffer (50mM Tris-HCl pH 7.5, 150mM NaCl, 0.25% sodium deoxycholate, 1% NP-40) supplemented with protease inhibitor cocktail and phosphatase inhibitor cocktail. 5 to 50μg of protein lysate were separated on 10% SDS-PAGE and transferred to polyvinylidene fluoride membrane. Membranes were blocked in 5% nonfat dry milk diluted in tris buffered saline with tween 20 (50mM Tris, 150mM NaCl, 0.05% Tween 20) and probed using the antibodies overnight at 4°C. GAPDH was used as loading control. Incubation with appropriate secondary HRP-labelled antibody (less than 1h) was followed by detection with Immobilon western chemiluminescent HRP substrate (Merck Millipore, Darmstadt, Germany).

### Immunoprecipitation and Mass Spectrometry analyses

Protein were extracted from HEK293 cells overexpressing UPF3B_L (ENST00000276201) or UPF3B_S (ENST00000636792) and subjected to immunoprecipitation with Pierce Anti-HA Magnetic Beads (Thermofisher 88836) according to manufacturer protocol. Immunoprecipitations were validated by WB as previoulsy done (33) before the mass spectrometry analyses (Proteomic platform, IGBMC). Briefly, samples were treated with LysC/Trypsine for liquid digestion and injected in Orbitrap ELITE / C18 Accucore 50cm (20μL 0.1%TFA / 1μf) for 2h runs in triplicate. Data were processed with Proteome Discoverer 2.2 software using Homosapiens_190716_reviewed. fasta and contaminants_190528.fasta databases. To consider a protein as a candidate interactor, we applied the thresholds to keep only protein with 1) at least two unique peptides, 2) a positive value of Peptide-Spectrum Matching (PSM) in each replicate 3) a positive eXtracted Ion Chromatogram (XIC) value in at least one replicate, 4) a sum of PSM for the three replicates of the control (empty) condition inferior to 5, 5) a sum of PSM for the three replicates of the test condition (UPF3B_L or UPF3B_L) greater or equal to 5 and 6) a ratio between the Normalized Spectral Abundance Factor (NSAF) of the test condition (UPF3B_L or UPF3B_L) and the NSAF of the control (empty) condition greater or equal to 2. Enrichment analysis were realized using DAVID (Database for Annotation, Visualization and Integrated Discovery) or the GO (Gene Ontology Resource).

## RESULTS

### Identification of genes differentially expressed in individuals with Renpenning syndrome

In order to identify the effect of PQBP1 deficiency on the regulation of gene expression and alternative splicing, we performed sequencing of mRNA extracted from lymphoblastoid cell lines (LCL) obtained from unrelated control males (n=3) and males carrying pathogenic or likely pathogenic variants in *PQBP1* (n=5) (**Figure 1**, **Figure S1-S2-S3, Table S1**), including one patient with the recurrent truncating p.Arg153fs variant as well as four patients with missense variants p.Tyr65Cys, p.Arg243Trp and p.Pro244Leu variants (4,24,25,34) reported to affect PQBP1’s ability to interact with partners (10,23,34). Analysis of gene expression revealed 79 protein-coding genes differentially expressed (DEG) in individuals with variants in *PQBP1*, 53 up-regulated and 26 down-regulated (**Figure S3, Table S3**). We observed for instance a strong increase in *APP* expression in individuals with a pathogenic variant in *PQBP1*. RT-qPCR was used to confirm the results obtained by RNAseq on mRNA from the same individuals and from a second set of LCLs including additional: control individuals (n=3), individuals with *PQBP1* frameshift variants (n=2), individuals with *PQBP1* variant of unknown significance (VUS, n=3) and individual with a neurodevelopmental condition from another genetic origin, the Fragile-X syndrome (FXS, n=4). However, while the increase observed in patients has been confirmed by RT-qPCR on the first set of LCLs at the mRNA level (**Figure S3C**) and at the protein level (**Figure S3D**), this was failed to be confirmed by the analysis of the second set of LCL (**Figure S3E**), which detected a large variability in *APP* expression, regardless of the status of the individuals. Similar results were obtained for other genes found upregulated in the RNASeq analysis (**data not shown**). Transcriptomic analysis performed at exon level using DEXseq identified 117 different expressed exons (DEE) belonging to 97 protein-coding genes (**Table S4**). Among them, four exons corresponding to a non-canonical short isoform of the *UPF3B* gene (ENST00000636792, *UPF3B_S)*, were significantly more covered in patients (log2FC: 1.06-2.17, adjusted p-value: 1.4E-3 – 1.78E-5) (**Figure 2A, Figure S4A**). The increase was confirmed by RT-qPCR in the same individuals, even if it seems less pronounced for the individual with the p.Tyr65Cys (P4) than for the other patients, and also found significant in the second set of patients while not detected for individuals with FXS (**Figure 2B-C**). This effect was specific to the short isoform as no change was observed in RNAseq or RT-qPCR for the long *UPF3B* isoform (ENST00000276201, *UPF3B_L)* (**Figure S4B)**. To confirm that *PQBP1* loss leads to an upregulation of *UPF3B_S*, we treated two human Neural Stem Cell lines (hNSC), which are self-renewal homogenous precursors of cortical neurons, with a pool of siRNA directed against *PQBP1*. A knock-down (KD) of 40% of PQBP1 expression was obtained (**Figure 3A-B**). A significant increase of *UPF3B_S* but not *UPF3B_L* expression was identified in the KD condition (while no change was observed using a nonspecific siRNA, scramble) (**Figure 3C**), confirming what was observed in patients with pathogenic variants in *PQBP1*. This increase was confirmed in various cell types after *PQBP1*-KD (HeLa, fibroblasts, HEK293, etc)(data not shown).

**Figure 1.**
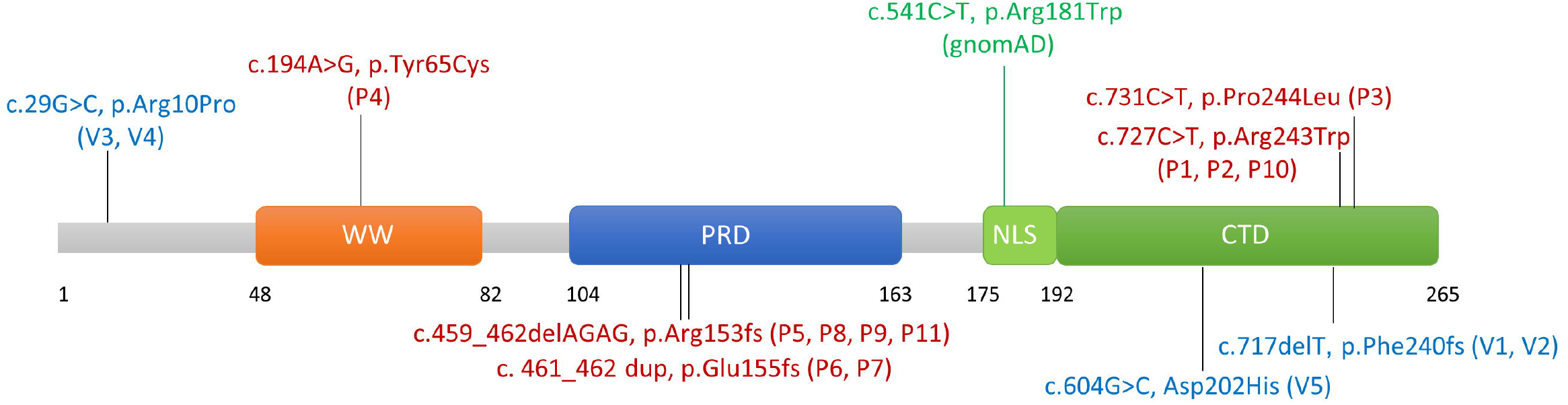
Variants in *PQBP1* in individuals with ID. Schematic representation of PQBP1 protein with its different domains (WW: WW domain, PRD: Proline-rich domain, NLS: nuclear localization signal, CTD:C-terminal domain) and the different variants analyzed in this study: on the top, missense variants; on the bottom, truncating variants; in green, missense variant found in males from gnomAD population and considered therefore as non-disease causing; in red, variants reported as disease causing or likely disease-causing; in blue, variants of uncertain significance.

**Figure 2.**
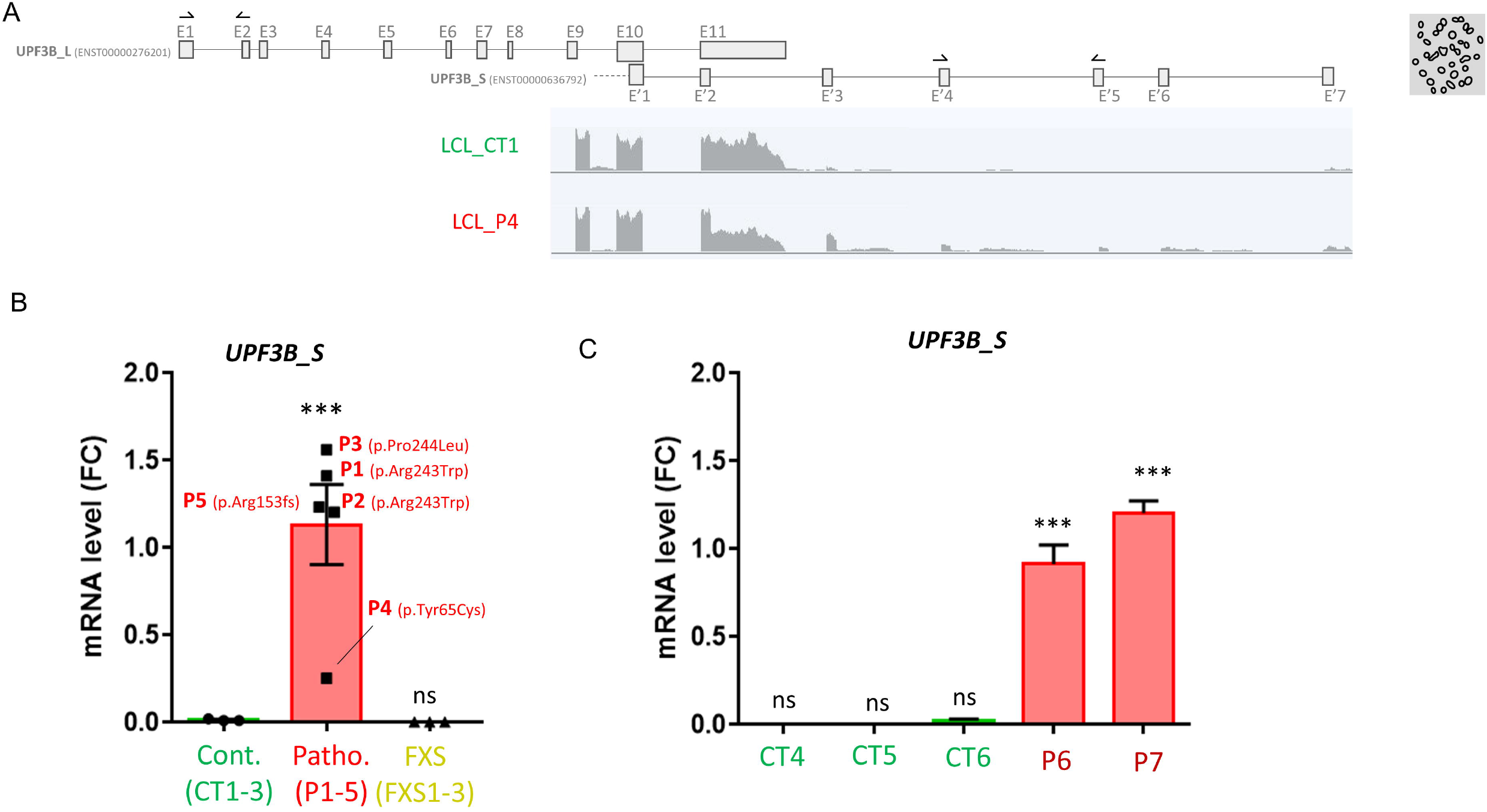
Transcriptomic study in LCL from individuals with pathogenic variants in *PQBP1* reveals an increase in the expression of a non-canonical isoform of *UPF3B*. (**A**) Schematic representation of the different exons composing the canonical longest isoform of *UPF3B* (ENST00000276201, *UPF3B_L)* as well as the shorter 3’ isoform (ENST00000636792, *UPF3B_S)*. IGV visualization of RNAseq data from one control individual (CT1) and one patient (P4). qPCR primers used for RT-qPCR validation are represented by black arrows. RT-qPCR analysis of *UPF3B_S* (normalized on *GAPDH* and *YWHAZ*) in (**B**) the first serie of LCL, control individuals (Cont, CT1-3, in green), individuals with pathogenic variants in *PQBP1* (Patho, in red) and individuals affected with Fragile-X syndrome (FXS, in yellow) (n ≥2 per cell line) and (**C**) in a second set of LCL from control individuals (CT4-6 in green) and individuals with the same frameshift p.Glu155fs variant in *PQBP1* (P6-7 in red).

**Figure 3.**
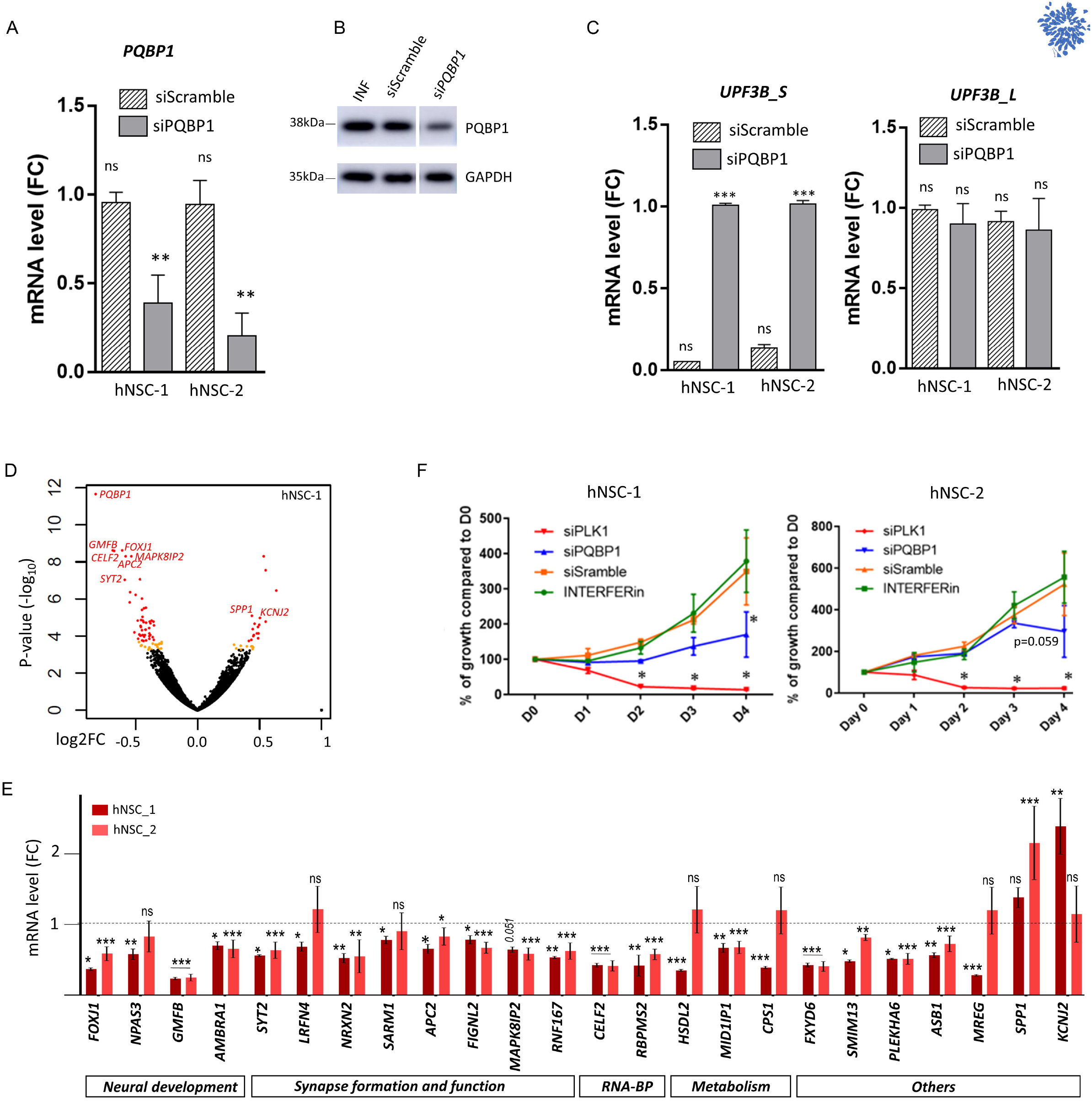
Consequences of *PQBP1* Knock-Down (KD) in human neural stem cells (hNSC) **(A)** RT-qPCR analysis of *PQBP1* mRNA (normalized on *GAPDH* and *YWHAZ)* in hNSC lines (hNSC-1 and hNSC-2) after 48h of transient knock-down (KD) of *PQBP1* using *siPQBP1* (n=3 per line). **(B)** Level of PQBP1 protein extracted from hNSC-1 after 48h transient *PQBP1*-KD (n=3 in total) **(C)** RT-qPCR analysis of *UPF3B_S* and *UPF3B_L* mRNA (normalized on *GAPDH* and *YWHAZ)* in hNSC-1 and hNSC-2 after a transient *PQBP1*-KD (n=3 per line). Multiple comparisons tests were performed using one-way ANOVA test with Dunnet’s correction: ns: not significant; ***: p-value <0.001; Errors bars represent SEM (standard error of the mean **(D)** Volcano plot showing RNA sequencing data (*siPQBP1* vs. INTERFERin treated hNSC-1). Genes with no significant change in expression are shown in black; genes deregulated with an adjusted p-value between 0.1 and 0.05 are shown in orange while genes significantly top deregulated (adjusted p-value <0.05, DEG) are shown in red. (**E**) RT-qPCR analysis of 24 DEG (normalized on *GAPDH* and *YWHAZ)* after *PQBP1*-KD in hNSC-1 (third serie) and in hNSC-2 (3 independent series in triplicates). Multiple comparison tests were performed using one-way ANOVA test with Dunnet’s correction: ns: not significant; *: p-value <0.01; **: p-value<0.01 ***: p-value <0.001; Errors bars represent SEM (**F**) Proliferation assay performed on hNSC-1 and hNSC-2 treated with lipofectant alone (INTERFERin) or transfected with *Scramble* siRNA (siScramble), *PQBP1* siRNA (siPQBP1) or *PLK1* siRNA (siPLK1), a control leading to cell apoptosis. At each time point (day 0, 1, 2, 3, and 4), cells were counted and data normalized to INTERFERin treatment (n=3 per line). Student’s t test comparison was done comparing to INTERFERin: *: p-value<0.05; Errors bars represent SEM

### Consequences of *PQBP1* knock-down in human neural stem cells (hNSC)

In order to identify additional DEG or DEE after *PQBP1*-KD in hNSC, we performed RNAseq of mRNA extracted from the first hNSC line (hNSC-1, in duplicates) (**Figure S2)**. Transcriptomic analysis revealed 59 significant DEG (12 up-regulated and 47 down-regulated) in *PQBP1*-KD cells compared to cells treated with transfection agent only (INTERFERin), while no gene was found significantly DE in the scramble siRNA condition (**Table S5, Figure 3D**). Unsurprisingly, the most significant DE gene was *PQBP1* (log2FC=−0.82, adjusted p-value =3.7E-8). To confirm these changes in gene expression, RT-qPCR was performed for the best candidate genes on a third series of hNSC-1 (n=3 samples per condition) as well as on three series of hNSC-2 (n=9 samples per condition in total) (**Figure 3E**). Among others, we identified that *PQBP1-KD* leads to a decrease in expression of genes encoding proteins essential for brain development, including transcription factors (*FOXJ1, NPAS3)*, neurotrophic factor (*GFMB)* or autophagic proteins (*AMBRA1*). A decrease of genes encoding proteins involved in neurite outgrowth and synaptic formation (*LRFN4, APC2* or *FIGNL2)* or synaptic function (*SYT2, NRXN2, SARM1, MAPK8IP2* or *RNF167)* was also observed. Two RNA-binding proteins involved in mRNA metabolism and particularly regulation of alternative splicing (*CELF2* and *RBPMS2*) were found to be down-regulated after *PQBP1*-KD. Decrease in expression of genes encoding metabolic enzymes, and especially enzymes involved in lipid or steroid metabolism (*HSDL2, MID1IP1)* have also been observed. As PQBP1 is known to play a role of cell cycle regulator in neural progenitors (18), which is consistent with the microcephaly observed in individuals affected by Renpenning syndrome, we decided to test if *PQBP1*-KD affect the proliferation of hNSC. We observed a significant decrease in hNSC proliferation 96 hours after *PQBP1*-KD in hNSC-1 when compared to transfection agent only (INTERFERin) or Scramble siRNA (**Figure 3F**). A similar tendency was obtained for hNSC-2 (p=0.059). Despite the described role of PQBP1 in splicing regulation, difference in exon coverage were observed in a few genes only using DEXSeq (n=3) (**Table S6**), but could not be confirmed by RT-PCR.

### The increase of *UPF3B_S* expression induced by *PQBP1* deficiency does not affect NMD

The *UPF3B_L* isoform encodes a protein involved in mRNA nonsense mediated decay (NMD), but the role of *UPF3B_S* is currently not known. This isoform is particularly expressed in testis and in EBV-immortalized lymphocytes but is also detected at lower level in other tissues such as cerebellum (GTex database). Therefore, to test if this increase in *UPF3B_S* expression caused by *PQBP1* deficiency affects NMD, we first searched if known NMD target genes (35) were DE in LCL from individuals with Renpenning syndrome or in *PQBP1*-KD hNSC but found no significant overlap. Finally, we proceeded to *PQBP1*-KD in fibroblasts carrying a truncating variant in the *DYRK1A* gene (c.1232dup, p.Arg413fs) known to be degraded by the NMD system (36). We observed an increase *of UPF3B_S* expression (2,4 and 6 days after inactivation) but with no effect on the *DYRK1A* mutant transcript, confirming that *PQBP1* inactivation has no obvious effect on NMD (data not shown). To study the role of UPF3B_S, we generated plasmids containing human *UPF3B_L* (ENST00000276201) and *UPF3B_S* cDNA sequences (ENST00000636792) and performed immunoprecipitation-coupled mass spectrometry on proteins extracted from HEK293 cells transfected with each of the constructs. We identified a total of 101 proteins: 13 proteins interacting with both isoforms, 74 proteins interacting preferentially or exclusively with UPF3B_L and only 14 preferentially or exclusively with UPF3B_S (**Table S7**). We retrieved eight of the known UPF3B interactors annotated in the STRING and BioGRID databases (**Figure 4**). As expected, we found that UPF1, UPF2 and ERF3A/GSPT1 interact only with UPF3B_L and not UPF3B_S, which lacks the interaction domain (**Figure 4**), while the EJC binding protein RBM8A (alias Y14) interacts with both isoforms. Among the proteins interacting preferentially with UPF3B_S, we identified proteins involved in antibacterial humoral response (LTF, SEMG1, IGHA1) and in redox homeostasis (PRDX4, SDHA).

**Figure 4.**
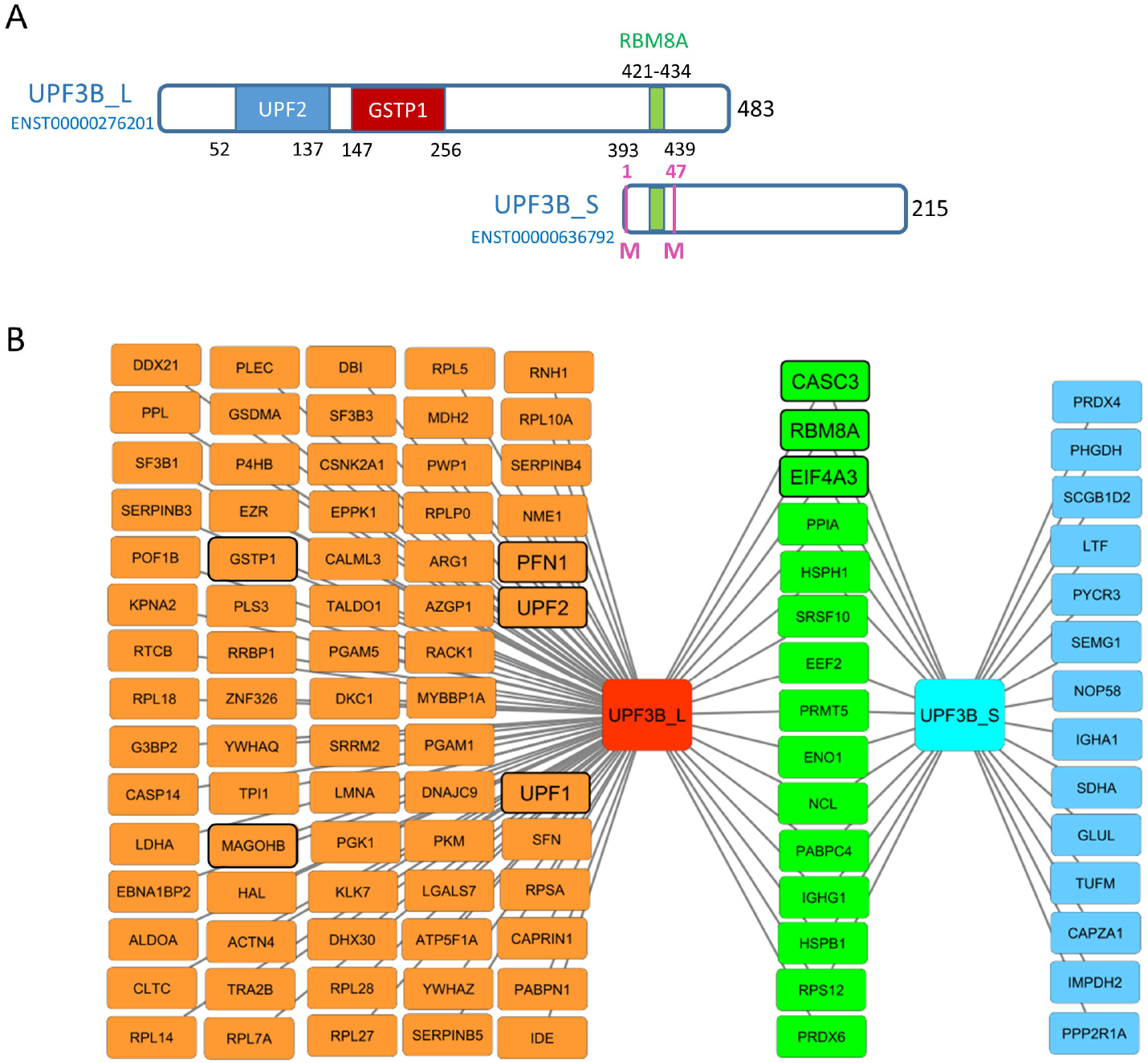
Representation and protein interactomes of UPF3B protein isoforms. (**A**) Schematic representation of UPF3B_L and UPF3B_S proteins with their domains known to interact with UPF2, GSTP1 and RBM8A proteins. (**B**) Protein interactome of UPF3B_L and UPF3B_S isoforms characterized by immunoprecipitation coupled to mass spectrometry (IP-MS) in HEK293 cells. In orange: interactors specific to UPF3B_L; in blue: interactors specific to UPF3B_S; in green: interactors of both UPF3B_L and UPF3B_S isoforms. Protein already described as interacting with UPF3B are circled with a black line.

### Increase in *UPF3B_S* expression is useful for variant testing

As the increase of *UPF3B_S* expression was observed in both cells from individuals with Renpenning syndrome and in all types of cells KD for *PQBP1* (**Figure 3C**), we intent to use it as a biomarker of *PQPB1* deficiency in order to test the functional consequences of variants of uncertain significance in this gene. We tested *UPF3B_S* expression in LCLs from individuals with *PQBP1* variants. We observed a significant increase of *UPF3B_S* in LCL obtained from two brothers carrying a distal frameshift variant, p.Phe240fs, leading to a protein 10 amino acids longer than the wild-type and previously reported by Hu et al. (24)(V1, V2). On the contrary, no significant increase was found in the individual carrying the p.Arg10Pro (V3)(**Figure 5A**). To avoid the immortalization step necessary to generate a LCL which is not routinely practiced by hospital laboratories, we tested if *UPF3B_S* expression could be detected on mRNA directly extracted form patients’ blood (PaxGene). As expected, no expression was detected in control individuals and individuals with FXS, while *UPF3B_S* was found expressed in blood from three individuals with pathogenic variants in *PQBP1* (P9-P11) (**Figure 5B**). Confirming the results obtained in LCLs, a slight but still not significant increase of *UPF3B_S* was observed in blood mRNA from individual V3 with the p.Arg10Pro variant.

**Figure 5.**
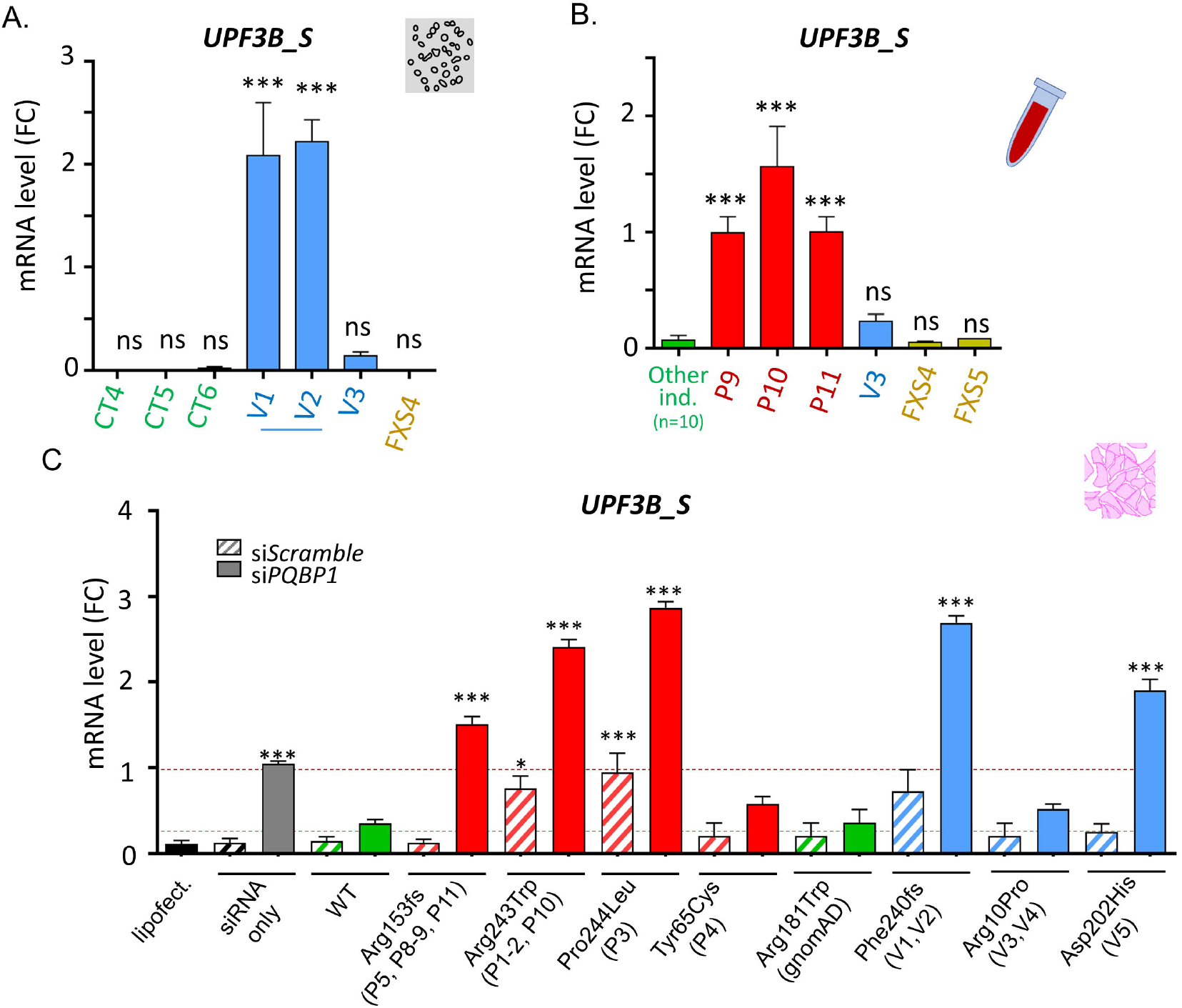
*UPF3B_S* increase is a robust marker of *PQBP1* loss of function and can be used for VUS testing. **(A)** RT-qPCR analysis of *UPF3B_S* (normalized on *GAPDH* and *YWHAZ)* in LCL from control individuals (CT4-6, in green), individuals with VUS in *PQBP1* (in blue, the two brothers V1 and V2 carrying p.Phe240fs and individual V3 carrying p.Arg10Pro) and in one individual affected from Fragile-X syndrome (FXS4, in yellow) (n=3 per cell line). LCL from individuals with pathogenic variants from the first set (P1-5) were used a calibrator. **(B)** RT-qPCR analysis of *UPF3B_S* (normalized on *GAPDH* and *YWHAZ)* in blood RNA samples (Paxgene) from individuals with ID without pathogenic variant in *PQBP1* (in green), individuals with pathogenic variants in *PQBP1* in red p.Arg153fs (P9, P11) and p.Arg243Trp (P10), individual with VUS in *PQBP1* (in blue) p.Arg10Pro (V3), and individuals affected from Fragile-X syndrome (FXS4, FSX5 in yellow) (n=1 per Paxgene) **(C)** RT-qPCR analysis of *UPF3B_S* mRNA (normalized on *GAPDH* and *YWHAZ*) in HeLa cells co-transfected by scramble or *PQBP1* siRNA and overexpressing WT or mutant PQBP1. Multiple comparisons tests were performed using one-way ANOVA test with Dunnet’s correction: ns: not significant; **: p-value <0.01; ***: p-value <0.001; Errors bars represent SEM.

In order to set up a functional assay which does not require patient’s material, we performed a complementation test in HeLa cells by co-transfecting *PQBP1* siRNA with optimized siRNA-resistant plasmids containing the wild-type or mutant human *PQBP1* cDNA (**Figure 5C, Figure S5**). We confirmed that *PQBP1-KD* increases *UPF3B_S* expression, while cotransfection with optimized WT *PQBP1* reverts this increase. As expected, transfection with the *PQBP1* containing a variant reported in two males from the GnomAD population, p.Arg181Trp, gave results similar to WT. On the contrary, co-transfection with mutant Arg153fs* *PQBP1* failed to revert the *UPF3B_S* increase, confirming the functional effect of this variant on *UPF3B_S*. Similarly, no complementation was obtained with p.Arg243Trp, p.Pro244Leu and p.Phe240fs mutants. and (**Figure 5C**). Interestingly, the *UPF3B_S* increase was even more pronounced for these mutants than for p.Arg153fs* and an increase was observed even without *PQBP1-KD*, suggesting that these mutants have a stronger effect than just a loss of *PQBP1*. The increase was not significant for the p.Tyr65Cys and p.Arg10Pro variants, confirming what we previously observed in LCL (**Figure 2B**) and suggesting that *UPF3B_S* increase is more pronounced for variants affecting the CTD domain of PQBP1 (truncating variants and missense variants in this domain) than for variants located in the N-terminal part of the protein. In addition, we used this complementation assay to test the effect of an additional variant located in this CTD and reported in literature in an individual with NDD, p.Asp202His (Morgan et al., 2015), and found a significant increase of *UPF3B_S* expression, suggesting a likely pathogenic effect (**Figure 5C**).

## DISCUSSION

In this study, we identified genes and exons differentially expressed in individuals with pathogenic variants in *PQBP1* or after a transient knock-down of *PQBP1* in human neural stem cells (hNSC). If we observed a high variability in gene expression in LCL, which prevented us to confirm several of the deregulations identified by RNA sequencing such as *APP* increase, the changes in gene expression observed in hNSC were robust and most of them were successfully replicated in a second independent hNSC line. Decrease in the expression of genes involved in brain development and synapse formation or in lipid metabolism were observed, which are consistent with the roles of PQBP1 previously described by others. For instance, a role of PQBP1 in regulation of dendrite length and branching has been described in mouse embryonic primary cortical neurons (12). PQBP1 was also found to regulate lipid metabolism: a decrease of the lipid content was observed in nematode intestinal cells, which could be related to the low body mass index observed in patients (37). Among the genes deregulated after *PQBP1*-KD, we found genes involved in other neurodevelopmental disorders such as *CELF2* (expression decreased after *PQBP1*-KD), an RNA binding protein mutated in a neurodevelopmental syndrome with ID and epilepsy (38), or *UPF3B* (expression increased for its non-canonical isoform *UPF3B_S*), another X-linked gene involved in ID (39). This increase of *UPF3B_S* was observed in different cell types after *PQBP1*-KD (hNSC, HeLa, fibroblasts, etc) and was also detected from patients’ material (LCL and blood). Compared to the canonical UPF3B_L transcript, this isoform involves an alternative transcription initiation site as well as an alternative splicing of exon 11 (the last exon of the canonical isoform).

There is no report about the biological function of this short isoform of *UPF3B*. *UPF3B_S* mRNA appears to be low expressed, except in testis, according to GTEx database. It is noticeable that genital manifestations, and especially microorchidia, were reported for individuals with Renpenning syndrome. A mild expression in Epstein Barr Virus-immortalized lymphoblastoid cells is also reported in GTEx, and might explain why we particularly detected this increase in LCL. If no expression of *UPF3B_S* is reported in GTEx in adult brain tissue, we cannot exclude that it could be expressed in a transient way during brain development. UPF3B_S predicted protein sequence contains motifs known to mediate interaction with RBM8A (alias Y14) but not those mediating interaction with UPF2/UPF1. Using immunoprecipitation coupled to mass spectrometry (IP-MS), we confirmed that UPF3B_S can bind to the EJC-protein RBM8A but fails to interact with NMD effectors UPF1 and UPF2. We could speculate that UPF3B_S could therefore exert a competitive antagonist activity to UPF3B_L on NMD. However, we did not observe NMD dysfunction after *PQBP*-KD in fibroblasts carrying a premature stop codon (PTC) and we observed no change in expression of natural target of NMD in hNSC or LCL. We found that UPF3B isoforms interact with proteins involved in mRNA binding and regulation of translation (ribosomal proteins and translation initiation or elongation factors). In particular, both isoforms interact with the translation elongation factor eEF2 (14), which is also known to be a partner of PQBP1. Indeed, PQBP1 binds to eEF2 and prevent its phosphorylation, which decreases translational elongation (14). Further experiments will be necessary to test whether UPF3B, known to play a role in the regulation of normal translation termination in addition to its role in NMD (40), could also be involved in regulation of translation elongation.

Among the few proteins detected as interacting specifically with UPF3B_S, some play a role in regulation of cell cycle (PRDX4, GLUL). In this study, we showed that *PQBP1*-KD affects the ability of hNSC to proliferate. This result is consistent with the microcephaly observed in individuals with Renpenning syndrome and with what was previously reported by other in non-human neuronal progenitors (18). Ito et al. observed, in nestin-Cre conditional KO mice for *Pqbp1*, which present with microcephaly, a loss of proliferation of neural progenitors probably due to an elongated cell cycle. As *Upf3b* KD in mouse neural progenitor cells leads on the contrary to an increase of cell proliferation (41), additional experiments would be needed to test if *UPF3B_S* expression increase might participates to the decrease of hNSC proliferation induced by *PQBP1*-KD. In addition, several other UPF3B_S-specific interactors are implicated in immune defense against bacterial infection (ex: LTF) while PQBP1 is known to play a role in response to infection as it can recognize viral DNA, interact with cGAS and LATS2 (42) and regulate immune response (16).

As *UPF3B_S* increase could be detected in blood RNA extracted from Renpenning patients, pathogenicity of future VUS identified in *PQBP1* could be evaluated from a simple blood RNA extraction. This biomarker seems to be more robust for variants located in the CTD domain than for variants located in the N-terminal part of the protein. To date, the CTD domain of PQBP1 is known to be involved in the interaction with the splicing factor TXNL4A alias U5-15kD (8) and it was recently shown that both Arg243Trp and Pro244Leu variants alter this binding (10,34). Tyr65Cys variant, which does not affect binding to TXNL4A (10), has a smaller effect on *UPF3B_S* expression increase. It is noticeable that some of these variants (Phe240fs, Pro244Leu Arg243Trp) tend to have a stronger effect than *PQBP1* deficiency and lead to *UPF3B_S* increase even when *PQBP1* WT is still expressed (siScramble condition, Figure 5C), suggesting a potential dominant-negative effect. We could therefore speculate that pathogenic variants in PQBP1 have different molecular effects depending of their location on the protein. However, it remains puzzling to note that no obvious phenotypic difference could be observed between patients with these different types of variants.

In conclusion, we describe here for the first time the consequences of *PQBP1* inactivation in human neural stem cells. We were able to show that cell proliferation is affected, as well as the expression of a about fifty genes, some of them known to be involved in other neurodevelopmental disorders. Despite the known role of PQBP1 in splicing regulation and the fact that the pathogenic variants are known to disrupt interactions with splicing factors (10,23), we failed to identify important changes in alternative splicing events. The only isoform-specific event we could detect was the increase of a non-canonical isoform of *UPF3B*, which has a still unknown function, but which can be used as a biomarker for *PQBP1* deficiency.

## DATA AVAILABILITY

Data have been submitted to Gene Expression Omnibus.

## ACKNOWLEDGEMENTS

The authors would like to thank the families for their participation and support. The authors also thank the Agence de Biomédecine, Fondation APLM for financial support. We also thank the GenomEast sequencing platform for performing RNASeq and for their help in data analysis, as well as Bastien Morlet from the Mass spectrometry platform and Anne Maglott from the high-throughput screening platform. We thank Camille Dourlens for her participation to the project as well as Josef Gecz, Frederic Laumonnier, Renaud Touraine and David Germanaud for scientific discussion and advice.

## ETHICS DECLARATION

Individuals were referred by clinical geneticists for genetic testing as part of routine clinical care. All patients enrolled and/or their legal representative have signed informed consent for research use and authorization for publication. IRB approval was obtained from the Ethics Committee of the Strasbourg University Hospital (CCPPRB) as well as from local institutions.

## WEB RESSOURCES

The URLs for online tools and data presented herein are:

CADD: https://cadd.gs.washington.edu/

ClinVar: http://www.ncbi.nlm.nih.gov/clinvar/

Clustal Omega: https://www.ebi.ac.uk/Tools/msa/clustalo/

Ensembl: https://www.ensembl.org/

GEO: https://www.ncbi.nlm.nih.gov/geo/

GnomAD:http://gnomad.broadinstitute.org/

GTEx: https://gtexportal.org/home/

Integrative Genomics Viewer (IGV): http://www.broadinstitute.org/igv/

OMIM: http://www.omim/org/

OrthoInspector database: https://www.lbgi.fr/orthoinspectorv3/databases

UCSC: http://genome.ucsc.edu/

## REFERENCES

1. Kalscheuer, V.M., Freude, K., Musante, L., Jensen, L.R., Yntema, H.G., Gécz, J., Sefiani, A., Hoffmann, K., Moser, B., Haas, S., et al. (2003) Mutations in the polyglutamine binding protein 1 gene cause X-linked mental retardation. Nat Genet, 35, 313–315.

2. Lenski, C., Abidi, F., Meindl, A., Gibson, A., Platzer, M., Frank Kooy, R., Lubs, H.A., Stevenson, R.E., Ramser, J. and Schwartz, C.E. (2004) Novel truncating mutations in the polyglutamine tract binding protein 1 gene (PQBP1) cause Renpenning syndrome and X-linked mental retardation in another family with microcephaly. Am J Hum Genet, 74, 777–780.

3. Kleefstra, T., Franken, C.E., Arens, Y.H.J.M., Ramakers, G.J.A., Yntema, H.G., Sistermans, E.A., Hulsmans, C.F.C.H., Nillesen, W.N., van Bokhoven, H., de Vries, B.B.A., et al. (2004) Genotype-phenotype studies in three families with mutations in the polyglutamine-binding protein 1 gene (PQBP1). Clin Genet, 66, 318–326.

4. Lubs, H., Abidi, F.E., Echeverri, R., Holloway, L., Meindl, A., Stevenson, R.E. and Schwartz, C.E. (2006) Golabi-Ito-Hall syndrome results from a missense mutation in the WW domain of the PQBP1 gene. J Med Genet, 43, e30.

5. Stevenson, R.E., Bennett, C.W., Abidi, F., Kleefstra, T., Porteous, M., Simensen, R.J., Lubs, H.A., Hamel, B.C.J. and Schwartz, C.E. (2005) Renpenning syndrome comes into focus. Am J Med Genet A, 134, 415–421.

6. Germanaud, D., Rossi, M., Bussy, G., Gérard, D., Hertz-Pannier, L., Blanchet, P., Dollfus, H., Giuliano, F., Bennouna-Greene, V., Sarda, P., et al. (2011) The Renpenning syndrome spectrum: new clinical insights supported by 13 new PQBP1-mutated males. Clin Genet, 79, 225–235.

7. Okazawa, H., Rich, T., Chang, A., Lin, X., Waragai, M., Kajikawa, M., Enokido, Y., Komuro, A., Kato, S., Shibata, M., et al. (2002) Interaction between mutant ataxin-1 and PQBP-1 affects transcription and cell death. Neuron, 34, 701–713.

8. Waragai, M., Junn, E., Kajikawa, M., Takeuchi, S., Kanazawa, I., Shibata, M., Mouradian, M.M. and Okazawa, H. (2000) PQBP-1/Npw38, a nuclear protein binding to the polyglutamine tract, interacts with U5-15kD/dim1p via the carboxyl-terminal domain. Biochem Biophys Res Commun, 273, 592–595.

9. Lee, B.J., Cansizoglu, A.E., Süel, K.E., Louis, T.H., Zhang, Z. and Chook, Y.M. (2006) Rules for nuclear localization sequence recognition by karyopherin beta 2. Cell, 126, 543–558.

10. Liu, X., Dou, L.-X., Han, J. and Zhang, Z.C. (2020) The Renpenning syndrome-associated protein PQBP1 facilitates the nuclear import of splicing factor TXNL4A through the karyopherin β2 receptor. J Biol Chem, 295, 4093–4100.

11. Waragai, M., Lammers, C.H., Takeuchi, S., Imafuku, I., Udagawa, Y., Kanazawa, I., Kawabata, M., Mouradian, M.M. and Okazawa, H. (1999) PQBP-1, a novel polyglutamine tract-binding protein, inhibits transcription activation by Brn-2 and affects cell survival. Hum Mol Genet, 8, 977–987.

12. Wang, Q., Moore, M.J., Adelmant, G., Marto, J.A. and Silver, P.A. (2013) PQBP1, a factor linked to intellectual disability, affects alternative splicing associated with neurite outgrowth. Genes Dev, 27, 615–626.

13. Kunde, S.A., Musante, L., Grimme, A., Fischer, U., Müller, E., Wanker, E.E. and Kalscheuer, V.M. (2011) The X-chromosome-linked intellectual disability protein PQBP1 is a component of neuronal RNA granules and regulates the appearance of stress granules. Hum Mol Genet, 20, 4916–4931.

14. Shen, Y., Zhang, Z.C., Cheng, S., Liu, A., Zuo, J., Xia, S., Liu, X., Liu, W., Jia, Z., Xie, W., et al. (2021) PQBP1 promotes translational elongation and regulates hippocampal mGluR-LTD by suppressing eEF2 phosphorylation. Mol Cell, 81, 1425–1438.e10.

15. Ikeuchi, Y., de la Torre-Ubieta, L., Matsuda, T., Steen, H., Okazawa, H. and Bonni, A. (2013) The XLID protein PQBP1 and the GTPase Dynamin 2 define a signaling link that orchestrates ciliary morphogenesis in postmitotic neurons. Cell Rep, 4, 879–889.

16. Yoh, S.M., Schneider, M., Seifried, J., Soonthornvacharin, S., Akleh, R.E., Olivieri, K.C., De Jesus, P.D., Ruan, C., de Castro, E., Ruiz, P.A., et al. (2015) PQBP1 Is a Proximal Sensor of the cGAS-Dependent Innate Response to HIV-1. Cell, 161, 1293–1305.

17. Ito, H., Yoshimura, N., Kurosawa, M., Ishii, S., Nukina, N. and Okazawa, H. (2009) Knock-down of PQBP1 impairs anxiety-related cognition in mouse. Hum Mol Genet, 18, 4239–4254.

18. Ito, H., Shiwaku, H., Yoshida, C., Homma, H., Luo, H., Chen, X., Fujita, K., Musante, L., Fischer, U., Frints, S.G.M., et al. (2015) In utero gene therapy rescues microcephaly caused by Pqbp1-hypofunction in neural stem progenitor cells. Mol Psychiatry, 20, 459–471.

19. Yang, S.-S., Ishida, T., Fujita, K., Nakai, Y., Ono, T. and Okazawa, H. (2020) PQBP1, an intellectual disability causative gene, affects bone development and growth. Biochem Biophys Res Commun, 523, 894–899.

20. Jensen, L.R., Chen, W., Moser, B., Lipkowitz, B., Schroeder, C., Musante, L., Tzschach, A., Kalscheuer, V.M., Meloni, I., Raynaud, M., et al. (2011) Hybridisation-based resequencing of 17 X-linked intellectual disability genes in 135 patients reveals novel mutations in ATRX, SLC6A8 and PQBP1. Eur J Hum Genet, 19, 717–720.

21. Abdel-Salam, G.M.H., Miyake, N., Abdel-Hamid, M.S., Sayed, I.S.M., Gadelhak, M.I., Ismail, S.I., Aglan, M.S., Afifi, H.H., Temtamy, S.A. and Matsumoto, N. (2018) Phenotypic and molecular insights into PQBP1-related intellectual disability. Am J Med Genet A, 176, 2446–2450.

22. Musante, L., Kunde, S.-A., Sulistio, T.O., Fischer, U., Grimme, A., Frints, S.G.M., Schwartz, C.E., Martínez, F., Romano, C., Ropers, H.-H., et al. (2010) Common pathological mutations in PQBP1 induce nonsense-mediated mRNA decay and enhance exclusion of the mutant exon. Hum Mutat, 31, 90–98.

23. Tapia, V.E., Nicolaescu, E., McDonald, C.B., Musi, V., Oka, T., Inayoshi, Y., Satteson, A.C., Mazack, V., Humbert, J., Gaffney, C.J., et al. (2010) Y65C missense mutation in the WW domain of the Golabi-Ito-Hall syndrome protein PQBP1 affects its binding activity and deregulates pre-mRNA splicing. J Biol Chem, 285, 19391–19401.

24. Hu, H., Haas, S.A., Chelly, J., Van Esch, H., Raynaud, M., de Brouwer, A.P.M., Weinert, S., Froyen, G., Frints, S.G.M., Laumonnier, F., et al. (2016) X-exome sequencing of 405 unresolved families identifies seven novel intellectual disability genes. Mol. Psychiatry, 21, 133–148.

25. Redin, C., Gérard, B., Lauer, J., Herenger, Y., Muller, J., Quartier, A., Masurel-Paulet, A., Willems, M., Lesca, G., El-Chehadeh, S., et al. (2014) Efficient strategy for the molecular diagnosis of intellectual disability using targeted high-throughput sequencing. J. Med. Genet., 51, 724–736.

26. Boissart, C., Nissan, X., Giraud-Triboult, K., Peschanski, M. and Benchoua, A. (2012) miR-125 potentiates early neural specification of human embryonic stem cells. Development, 139, 1247–1257.

27. Quartier, A., Chatrousse, L., Redin, C., Keime, C., Haumesser, N., Maglott-Roth, A., Brino, L., Le Gras, S., Benchoua, A., Mandel, J.-L., et al. (2018) Genes and Pathways Regulated by Androgens in Human Neural Cells, Potential Candidates for the Male Excess in Autism Spectrum Disorder. Biol. Psychiatry.

28. Kim, D., Pertea, G., Trapnell, C., Pimentel, H., Kelley, R. and Salzberg, S.L. (2013) TopHat2: accurate alignment of transcriptomes in the presence of insertions, deletions and gene fusions. Genome Biol., 14, R36.

29. Langmead, B. and Salzberg, S.L. (2012) Fast gapped-read alignment with Bowtie 2. Nat Methods, 9, 357–359.

30. Anders, S., Pyl, P.T. and Huber, W. (2015) HTSeq--a Python framework to work with high-throughput sequencing data. Bioinformatics, 31, 166–169.

31. Benjamini, Y Controlling the False Discovery Rate: A Practical and Powerful Approach to Multiple Testing. Controlling the False Discovery Rate: A Practical and Powerful Approach to Multiple Testing.

32. Anders, S., Reyes, A. and Huber, W. (2012) Detecting differential usage of exons from RNA-seq data. Genome Res, 22, 2008–2017.

33. Mattioli, F., Isidor, B., Abdul-Rahman, O., Gunter, A., Huang, L., Kumar, R., Beaulieu, C., Gecz, J., Innes, M., Mandel, J.-L., et al. (2019) Clinical and functional characterization of recurrent missense variants implicated in THOC6-related intellectual disability. Hum. Mol. Genet., 28, 952–960.

34. Lopez-Martín, S., Albert, J., Peña Vila-Belda, M.D.M., Liu, X., Zhang, Z.-C., Han, J., Jiménez de Domingo, A., Fernández-Mayoralas, D.M., Fernández-Perrone, A.L., Calleja-Pérez, B., et al. (2021) A mild clinical and neuropsychological phenotype of Renpenning syndrome: A new case report with a maternally inherited PQBP1 missense mutation. Appl Neuropsychol Child, 1–7.

35. Nickless, A., Bailis, J.M. and You, Z. (2017) Control of gene expression through the nonsense-mediated RNA decay pathway. Cell Biosci, 7, 26.

36. Courraud, J., Chater-Diehl, E., Durand, B., Vincent, M., del Mar Muniz Moreno, M., Boujelbene, I., Drouot, N., Genschik, L., Schaefer, E., Nizon, M., et al. (2021) Integrative approach to interpret DYRK1A variants, leading to a frequent neurodevelopmental disorder. medRxiv, 2021.01.20.21250155.

37. Takahashi, K., Yoshina, S., Masashi, M., Ito, W., Inoue, T., Shiwaku, H., Arai, H., Mitani, S. and Okazawa, H. (2009) Nematode homologue of PQBP1, a mental retardation causative gene, is involved in lipid metabolism. PLoS One, 4, e4104.

38. Itai, T., Hamanaka, K., Sasaki, K., Wagner, M., Kotzaeridou, U., Brösse, I., Ries, M., Kobayashi, Y., Tohyama, J., Kato, M., et al. (2021) De novo variants in CELF2 that disrupt the nuclear localization signal cause developmental and epileptic encephalopathy. Hum Mutat, 42, 66–76.

39. Tarpey, P.S., Raymond, F.L., Nguyen, L.S., Rodriguez, J., Hackett, A., Vandeleur, L., Smith, R., Shoubridge, C., Edkins, S., Stevens, C., et al. (2007) Mutations in UPF3B, a member of the nonsense-mediated mRNA decay complex, cause syndromic and nonsyndromic mental retardation. Nat Genet, 39, 1127–1133.

40. Deka, B., Chandra, P. and Singh, K.K. (2021) Functional roles of human Up-frameshift suppressor 3 (UPF3) proteins: From nonsense-mediated mRNA decay to neurodevelopmental disorders. Biochimie, 180, 10–22.

41. Jolly, L.A., Homan, C.C., Jacob, R., Barry, S. and Gecz, J. (2013) The UPF3B gene, implicated in intellectual disability, autism, ADHD and childhood onset schizophrenia regulates neural progenitor cell behaviour and neuronal outgrowth. Hum Mol Genet, 22, 4673–4687.

42. He, T.-S., Dang, L., Zhang, J., Zhang, J., Wang, G., Wang, E., Xia, H., Zhou, W., Wu, S. and Liu, X. (2021) The Hippo signaling component LATS2 enhances innate immunity to inhibit HIV-1 infection through PQBP1-cGAS pathway. Cell Death Differ.

