## Supplementaries for "Molecular consequences of *PQBP1* deficiency, involved in the X-linked Renpenning syndrome"

**Figure S1. Characterization of *PQBP1* variants in patients' LCL**

**(A)** Sanger sequencing performed to validate the presence of the different *PQBP1* variants in LCL lines. **(B)** Level of PQBP1 WT and mutant proteins extracted from control individuals (CT, in green), patients with pathogenic variants in *PQBP1* (P, in red) and patients with VUS in *PQBP1* (V, in blue).

**Figure S2. Experimental design of transcriptomic studies**

List of RNA-Sequencing and RT-qPCR analysis performed in human neural stem cells (hNSC: SA001 and GM01869), in lymphoblastoid cell lines (LCL) obtained from control individuals (CT), individuals with pathogenic variants in *PQBP1* (P), individuals with variants of uncertain significance (VUS) in *PQBP1* (V) or individuals with pathogenic variants in *FMR1* (FXS).

**Figure S3. Transcriptomic analysis performed in LCL from patients with pathogenic variants in *PQBP1* compared to control individuals**

**(A)** Principal Component Analysis of transcriptomic analysis performed on LCL from individuals with pathogenic variant in *PQBP1* (P1-5) vs control individuals (CT1-3) **(B)** Volcano plot showing changes in gene expression in individuals with pathogenic variant in *PQBP1* (P1-5). P-value (-log10) is expressed for each gene in function of log2fold change (log2FC). Genes not significantly deregulated are shown in black; while significantly deregulated genes (adjusted p-value <0.05) are shown in red. **(C)** RT-qPCR analysis of *APP* mRNA level (normalized on *GAPDH* and *YWHAZ*) in LCL from the first set: control individuals (Cont, in green), individuals with pathogenic variants in *PQBP1* (Patho, in red) and individuals affected by Fragile-X syndrome (FXS, in yellow) (n=1 per cell line) **(D)** Western Blot analysis on total protein extracts from LCL of individuals with *PQBP1* variants and control individuals using APP antibody and beta-Tubulin antibody as a control **(E)** RT-qPCR analysis of *APP* mRNA level (normalized on *GAPDH* and *YWHAZ*) in LCL from additional control individuals (CT4-6, in green), individuals with *PQBP1* pathogenic variants (P6-7, in red) or VUS (V1-3, in blue) and individual affected by Fragile-X syndrome (FSX4, in yellow) (n=3 per cell line); control LCL from the first set (CT 1-3) were used a calibrator. Multiple comparisons tests were performed using one-way ANOVA test with Dunnet's correction: ns: not significant; ***: adjusted p-value<0.001; Errors bars represent SEM (standard error of the mean).

#### Figure S4. Expression of *UPF3B* isoforms

**(A)** Visualization of DEXseq results of RNA-Sequencing performed in LCL, for the *UPF3B* locus. The normalized counts are represented for control individuals and individuals carrying *PQBP1* pathogenic variants. *: exons found significantly upregulated (adjusted p-value <0.05) in patients by DEXseq analysis. Genomic positions are given according to hg38 version of the genome. **(B)** RT-qPCR analysis of *UPF3B* long isoform (*UPF3B_L)* mRNA level (normalized on GAPDH and YWHAZ) in LCL from control individuals (CT, in green), individuals with pathogenic variants in *PQBP1* (Patho, in red) and individuals affected by Fragile-X syndrome (FXS, in yellow) (n =1 per cell line). Multiple comparisons tests were performed using one-way ANOVA test with Dunnet's correction: ns: not significant; Errors bars represent SEM (standard error of the mean).

#### Figure S5. Transfection with *PQBP1* siRNA in HeLa cells leads to a decrease of endogenous *PQBP1* expression.

RT-qPCR analysis of endogenous (endo) *PQBP1* mRNA using primers not amplifying *PQBP1* constructs in HeLa after co expressing si*PQBP1* and WT or mutant *PQBP1* cDNA sequences. Multiple comparisons tests were performed using onea-way ANOVA test with Dunnet's correction (each condition compared to "siScramble only" condition): ns: not significant; ***: adjusted p-value<0.001; Errors bars represent SEM (standard error of the mean).

### Supplementary tables

#### Sup Table 1. List of the samples and variants analyzed in this study

For each individual is indicated if some LCL was available, if they were included in the RNA-Seq analysis, if mRNA sample extracted from total blood (Paxgene) was available, if the variant was included in the functional complementation assay and, for missense variants, the prediction scores according to Polyphen2 and CADD.

**Sup Table 2. List of the primers used for RT-qPCR analysis**

**Sup Table 3. List of Differentially Expressed Genes (DEG) in LCL from individuals with pathogenic variants in *PQBP1* identified by DEseq analysis**

**Sup Table 4. List of Differentially Expressed Exons (DEE) in LCL from individuals with pathogenic variants in *PQBP1* identified by DEXseq analysis**

**Sup Table 5. List of Differentially Expressed Genes (DEG) in hNSC-1 after *PQBP1* knock-down (KD) identified by DEseq analysis**

**Sup Table 6. List of Differentially Expressed Exons (DEE) in hNSC-1 after *PQBP1* knock-down (KD) identified by DEXseq analysis**

**Sup Table 7. Protein interactors of UPF3B_S and UPF3B_L identified by Immunoprecipitation coupled to Mass Spectrometry**

PSM: Peptide-Spectrum Matching; NSAF: Normalized Spectral Abundance Factor

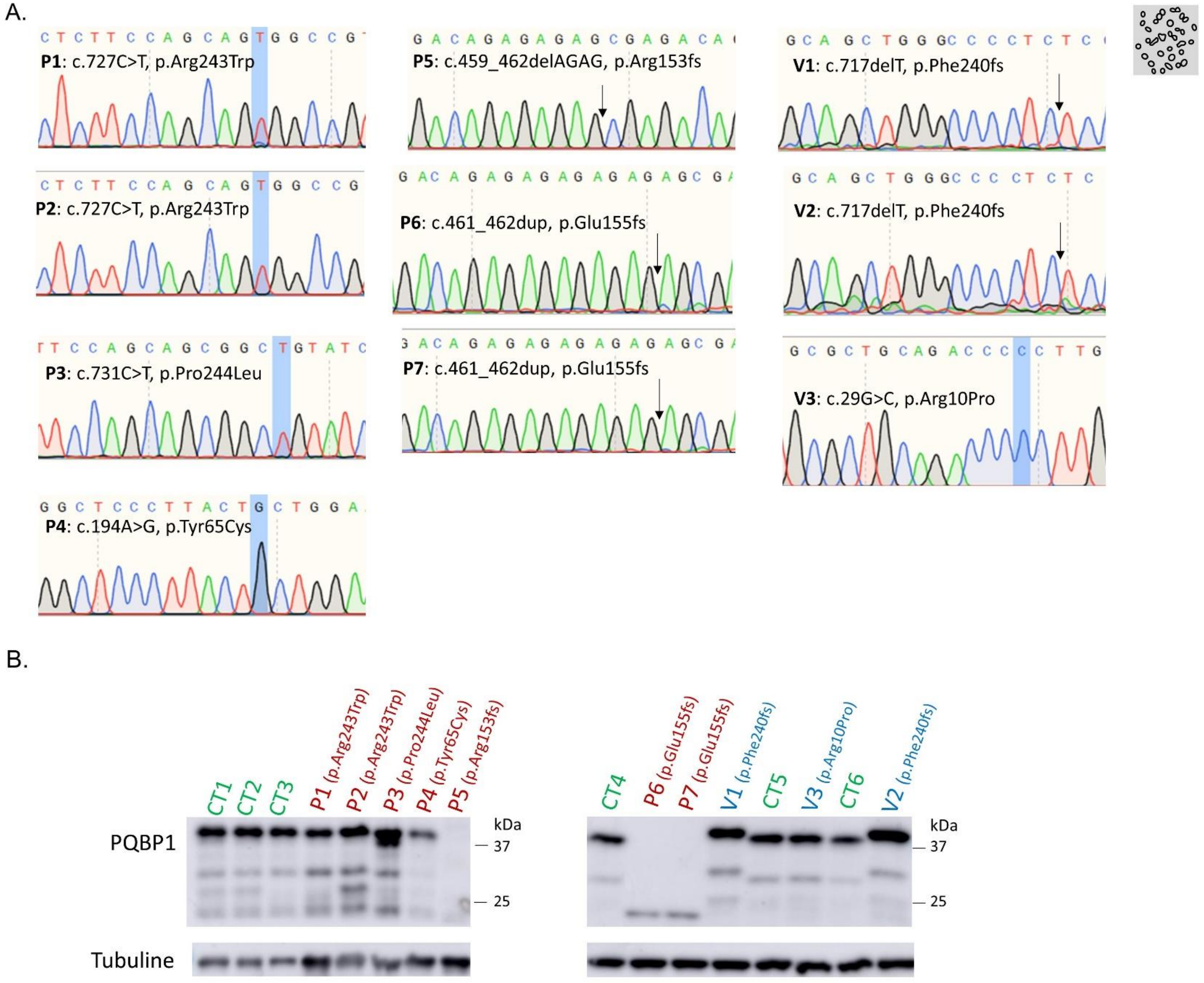
A.
P1: c.727C>T, p.Arg243Trp
P2: c.727C>T, p.Arg243Trp
P3: c.731C>T, p.Pro244Leu
P4: c.194A>G, p.Tyr65Cys
P5: c.459_462delAGAG, p.Arg153fs
P6: c.461_462dup, p.Glu155fs
P7: c.461_462dup, p.Glu155fs
V1: c.717delT, p.Phe240fs
V2: c.717delT, p.Phe240fs
V3: c.29G>C, p.Arg10Pro
B.
CT1
CT2
CT3
P1 (p.Arg243Trp)
P2 (p.Arg243Trp)
P3 (p.Pro244Leu)
P4 (p.Tyr65Cys)
P5 (p.Arg153fs)
CT4
P6 (p.Glu155fs)
P7 (p.Glu155fs)
V1 (p.Phe240fs)
CT5
V3 (p.Arg10Pro)
CT6
V2 (p.Phe240fs)
PQBP1
Tubuline
kDa
37
25

**Figure S1**

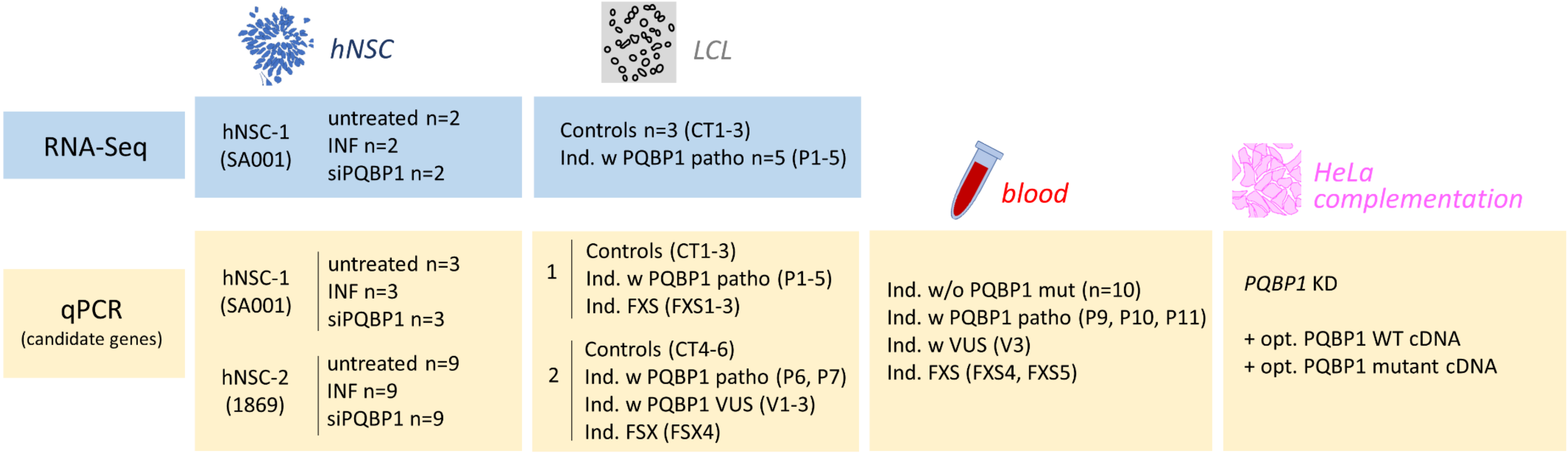
hNSC
LCL
RNA-Seq
hNSC-1
(SA001)
untreated n=2
INF n=2
siPQBP1 n=2
Controls n=3 (CT1-3)
Ind. w PQBP1 patho n=5 (P1-5)
blood
HeLa
complementation
qPCR
(candidate genes)
hNSC-1
(SA001)
untreated n=3
INF n=3
siPQBP1 n=3
hNSC-2
(1869)
untreated n=9
INF n=9
siPQBP1 n=9
1
Controls (CT1-3)
Ind. w PQBP1 patho (P1-5)
Ind. FXS (FXS1-3)
2
Controls (CT4-6)
Ind. w PQBP1 patho (P6, P7)
Ind. w PQBP1 VUS (V1-3)
Ind. FSX (FSX4)
Ind. w/o PQBP1 mut (n=10)
Ind. w PQBP1 patho (P9, P10, P11)
Ind. w VUS (V3)
Ind. FXS (FXS4, FXS5)
PQBP1 KD
+ opt. PQBP1 WT cDNA
+ opt. PQBP1 mutant cDNA

**Figure S2**

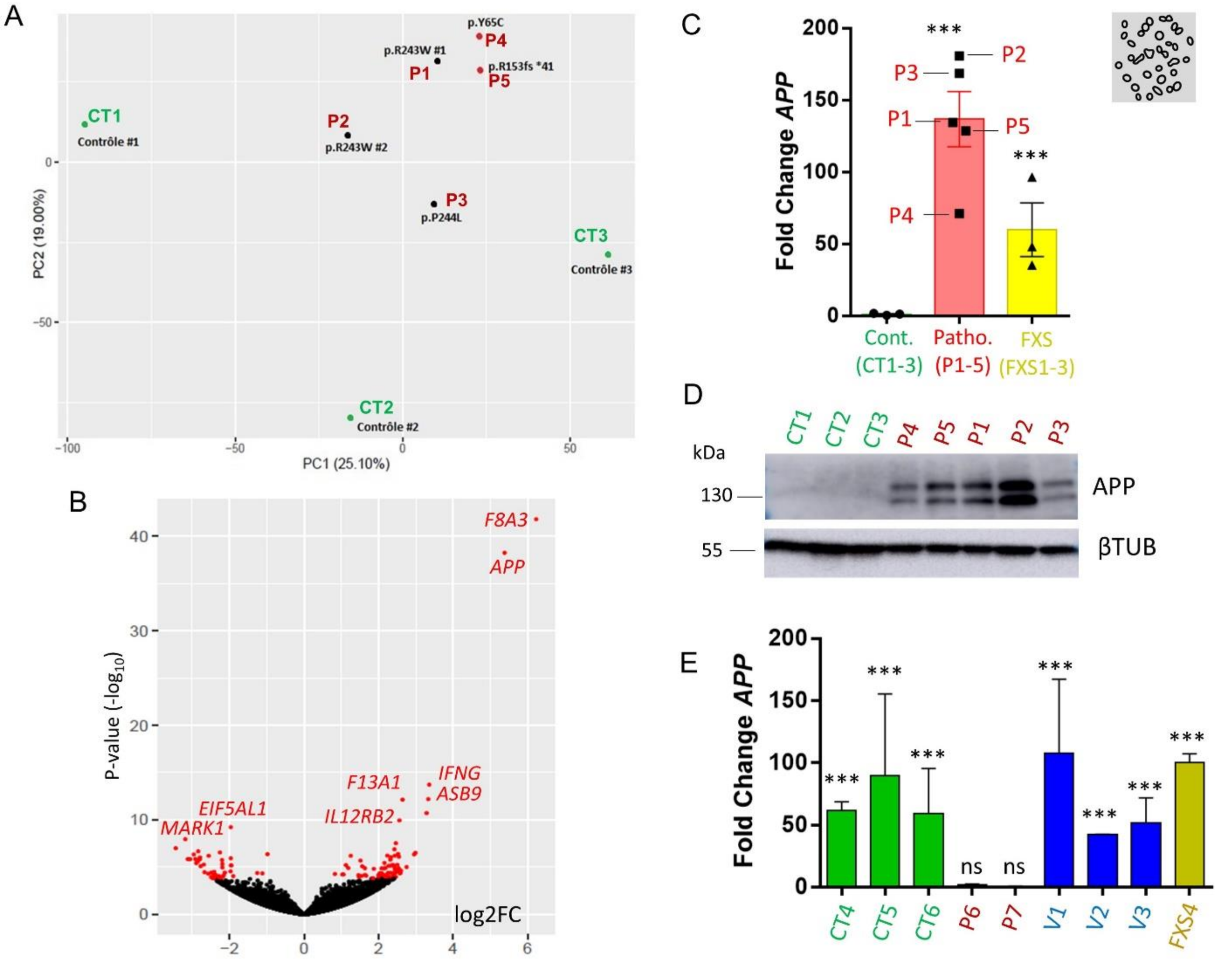
A
CT1
Contrôle #1
P2
p.R243W #2
p.R243W #1
P1
p.Y65C
P4
p.R153fs *41
P5
P3
p.P244L
CT3
Contrôle #3
CT2
Contrôle #2
PC2 (19.00%)
PC1 (25.10%)
B
F8A3
APP
IFNG
ASB9
F13A1
IL12RB2
EIF5AL1
MARK1
P-value (-log10)
log2FC
C
Fold Change APP
Cont. (CT1-3)
Patho. (P1-5)
FXS (FXS1-3)
D
CT1 CT2 CT3 P4 P5 P1 P2 P3
kDa
130
55
APP
βTUB
E
Fold Change APP
CT4 CT5 CT6 P6 P7 V1 V2 V3 FXS4
ns

**Figure S3**

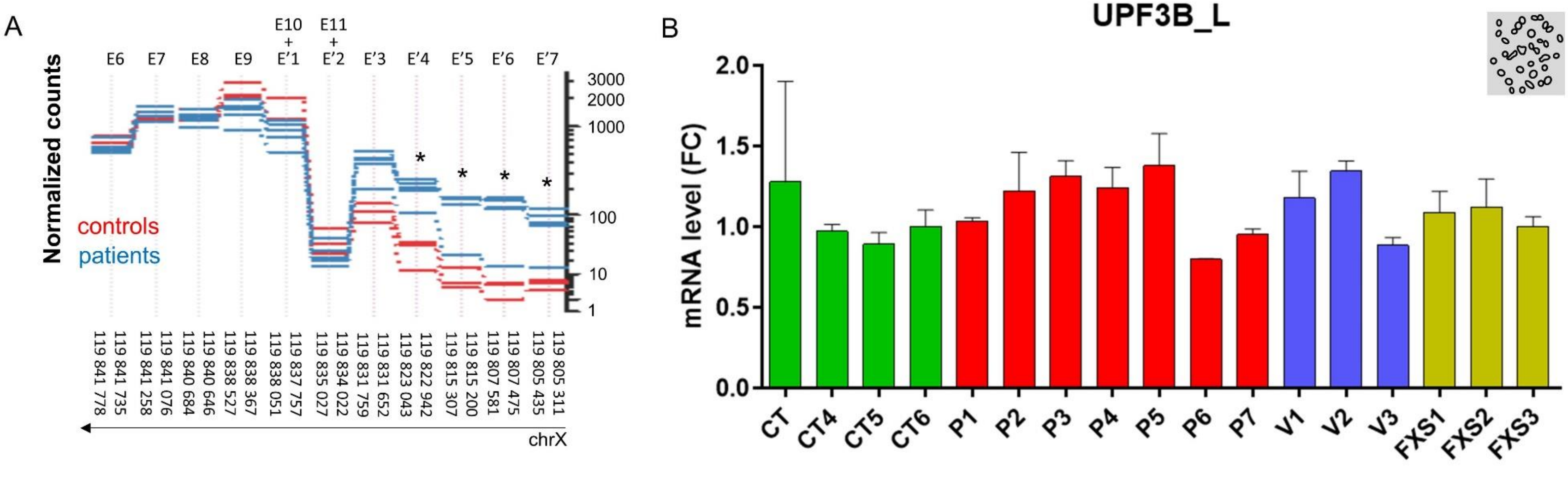
A
E6 E7 E8 E9 E10 + E'1 E11 + E'2 E'3 E'4 E'5 E'6 E'7
Normalized counts
controls
patients
3000
2000
1000
100
10
1
119 841 778
119 841 735
119 841 258
119 841 076
119 840 684
119 840 646
119 838 527
119 838 367
119 838 051
119 837 757
119 835 027
119 834 022
119 831 759
119 831 652
119 823 043
119 822 942
119 815 307
119 815 200
119 807 581
119 807 475
119 805 435
119 805 311
chrX
B
UPF3B_L
mRNA level (FC)
2.0
1.5
1.0
0.5
0.0
CT CT4 CT5 CT6 P1 P2 P3 P4 P5 P6 P7 V1 V2 V3 FXS1 FXS2 FXS3

**Figure S4**

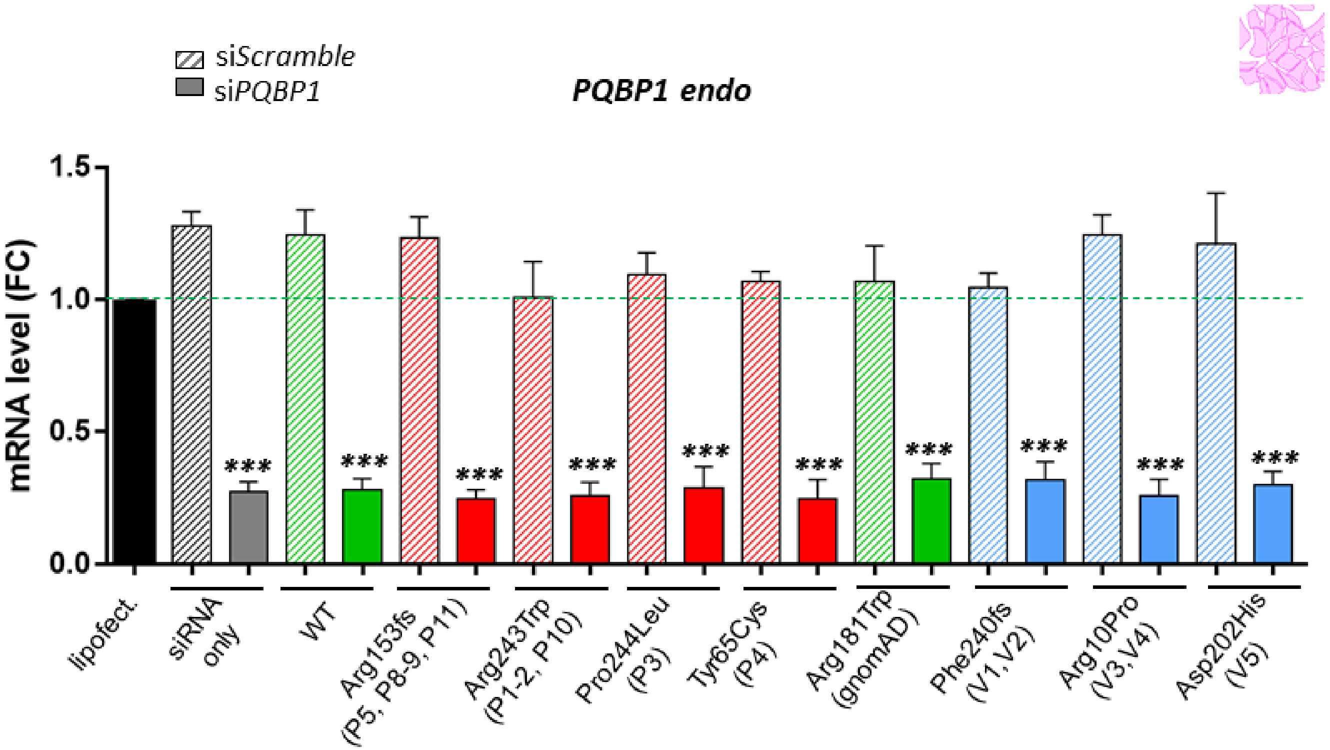
siScramble
siPQBP1
PQBP1 endo
mRNA level (FC)
1.5
1.0
0.5
0.0
***
lipofect.
siRNA only
WT
Arg153fs (P5, P8-9, P11)
Arg243Trp (P1-2, P10)
Pro244Leu (P3)
Tyr65Cys (P4)
Arg181Trp (gnomAD)
Phe240fs (V1,V2)
Arg10Pro (V3,V4)
Asp202His (V5)

**Figure S5**
